# Brain fingerprinting using fMRI spectral signatures on high-resolution cortical graphs

**DOI:** 10.1101/2023.03.14.532594

**Authors:** Carlo Ferritto, Maria Giulia Preti, Stefano Moia, Dimitri Van De Ville, Hamid Behjat

## Abstract

Resting-state fMRI has proven to entail subject-specific signatures that can serve as a fingerprint to identify individuals. Conventional methods are based on building a connectivity matrix based on correlation between the average time course of pairs of brain regions. This approach, first, disregards the exquisite spatial detail manifested by fMRI due to working on average regional activities, second, cannot disentangle correlations associated to cognitive activity and underlying noise, and third, does not account for cortical morphology that spatially constraints function. Here we propose a method to address these shortcomings via leveraging principles from graph signal processing. We build high spatial resolution cortical graphs that encode each individual’s cortical morphology and treat region-specific, whole-hemisphere fMRI maps as signals that reside on the graphs. fMRI graph signals are then decomposed using systems of graph spectral kernels to extract structure-informed functional signatures, which are in turn used for fingerprinting. Results on 100 subjects showed the overall superior subject differentiation power of the proposed signatures over the conventional method. Moreover, placement of the signatures within canonical functional brain networks revealed the greater contribution of high-level cognitive networks in subject identification.

## 1. INTRODUCTION

Functional Magnetic Resonance Imaging (fMRI) has revolutionised our understanding of brain function and has become a popular tool in cognitive neuroscience research. Recent studies have shown that fMRI data can be used to identify individuals based on their unique brain activity patterns, a technique known as “fingerprinting” [1], that is important e.g. to assess test-retest reliability of functional connectivity MRI [2]. Beneficial in part because of its simplicity, most fMRI fingerprinting methods have continued to rely on functional connectomes [3] (FC), i.e. subject-specific full graphs wherein vertices represent brain regions and edge weights represent the strength of statistical dependency between activity time-courses of pairs of brain regions (e.g. with Pearson’s correlation). Correlation based measures capture, however, not only couplings between neural activity but also underlying noise [4, 5]. As a remedy, FC denoising strategies based on learning a latent space embedding from a population of subjects have been proposed [6, 7]. More recently, alternative measures to FC have been leveraged, in particular, using graph learning to infer alternative functional graphs [8] or using principles from graph signal processing (GSP) [9] to derive structure-function coupling signatures [10, 11].

Aside form shortcomings related to correlation-based FC, fingerprinting strategies dominantly rely on coarsening high spatial resolution voxel-wise maps (100K+ elements) down to atlas-based, region-wise spatially averaged maps (1000 or less elements) [1, 6–8, 11]. Intrinsic organisation of average regional activities are thus the embedding to extract features, limiting in-depth exploitation of the high spatial resolution provided by fMRI. In this work we take advantage of the full spatial resolution of fMRI data, interpreting fMRI maps as functions residing on subject-specific, voxel-wise brain graphs [12– In particular, we show how GSP-based spectral signatures of fMRI data on cerebral hemisphere cortex (CHC) graphs [14, 16] can provide informative signatures for subject identification. We decompose resting-state fMRI data using systems of spectral heat kernels [13, 17], and, in turn, treat the energy retained in each filtered signal [16] as a feature. Our proposed method does indeed rely on using a brain parcellation [18], albeit not for coarsening cortical activity maps, but rather to derive voxel-wise cortical maps that are more informative than raw fMRI maps, each associated to a specific cortical region [19]. Thus, we are able to assess i) the change in performance relative to the choice of parcellation, and ii) the importance of different cortical regions for fingerprinting.

## 2. METHODS

### 2.1. Graph signal processing fundamentals

Let 𝒢 denote an undirected, unweighted, without self-loop graph with *N* vertices, characterized by an adjacency matrix **A** ∈ ℝ^*N×N*^ with elements *A*_*i,j*_ = 1 if an edge connects vertices *i* and *j*, and *A*_*i,j*_ = 0 otherwise. The graph symmetric normalized Laplacian matrix **L** is defined as 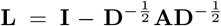, where **I** denotes the identity matrix, and **D** denotes the diagonal degree matrix with elements *D*_*i,i*_ = Σ_*j*_ *A*_*i,j*_. The eigendecomposition of **L** gives **L** = **UΛU**^*T*^, where **Λ** is a diagonal matrix storing the eigenvalues 0 = *λ*_1_ ≤ *λ*_2_ … ≤ *λ*_*N*_ ≤ 2 and **U** = [**u**_1_, **u**_2_, · · ·, **u**_*N*_] is a matrix which stores the associated eigenvectors **u**_*n*_, *n* = 1, …, *N* in its columns; the eigenvalues represent spatial frequencies and define the graph spectrum [20]. A graph filter can be conveniently defined in the spectral domain as a continuous kernel *h*(·) : [0, *λ*_*N*_] → ℝ, using which a graph signal **f** can be filtered, denoted **f**_*h*(*·*)_, by modulating its spectral representation and reconstruction back to vertex domain as [21]

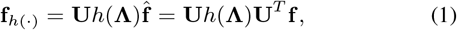

where *h*(**Λ**) is a diagonal matrix with its *n*-th entry being *h*(*λ*_*n*_). For large graphs, a graph signal can be filtered in a computationally efficient way via an approximation scheme, wherein a polynomial estimate of *h*(·), denoted *p*(·) : [0, *λ*_*N*_] → ℝ, can be used to compute the filtered signal in the vertex domain as:

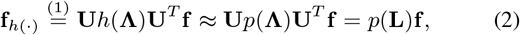

where the last equality is based on invoking the property **Lu**_*n*_ = *λ λ*_*n*_**u**_*n*_ → *p*(**L**)**u**_*n*_ = *p*(*λ*_*n*_)**u**_*n*_. Thus, graph signal filtering can be implemented via matrix operations on **L**, of the same order as the polynomial order, obviating the need to resort to the spectral representation of the signal. In the following, we only consider polynomial approximations of kernels, denoting such kernels as *p*(·), or its sub-indexed versions *p*_*k*_(·). For any given kernel *h*(·), we estimate the minimum required order for the approximated polynomial *p*(·) such that ∀*λ* ∈ [0, *λ*_max_], |*p*(*λ*) − *h*(*λ*) | ≤ 0.0001.

Graph signal filtering is shift-variant. In particular, given a filtering matrix **H** ∈ ℝ^*N×N*^ :

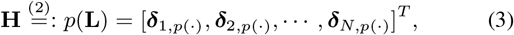

where ***δ***_*n,p*(*·*)_ ∈ ℝ^*N×*1^ (row *n* in **H**) represents the impulse response—a.k.a. graph atom—associated to *p*(·) when instantiated at vertex *n*; that is, filtering is performed by applying a shift-varinat impulse response set, each of which adapt to the local graph structure at their respective vertex. With this view, by virtue of the matched filter argument, **f**_*p*_[*n*] = **f** ^*T*^ ***δ***_*n,p*(*·*)_ is a quantification of the extent of signal energy associated to the spectral profile given by *p*(·) in the local vicinity of vertex *n*.

Given a system of spectral kernels (SOSKS):

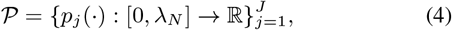

for which there exists bounds 0 *< B*_1_ *< B*_2_ *<* ∞ such that ∀*λ* ∈|0,*λ*_*N*_) 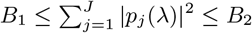, the dictionary of atoms

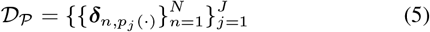

forms a frame for the Hilbert space of real valued graph signals, satisfying ∀**f** ∈ ℝ^*N×*1^, 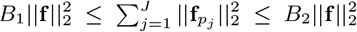. Using 𝒟_*P*_, an *N* dimensional graph signal **f** can be decomposed into a *JN* -dimensional embedding 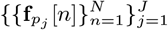, using which subtle signatures can be extracted from the signal. Alternatively, the dimensionality of the embedding can be reduced to *J* by considering filtered signal norms, thus, obtaining a feature set 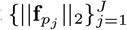. In this work we use the latter strategy due to the sheer size of the graphs, whereas the former option will be considered in future work.

### 2.2. Heat-SOSKS

In this work we focus on using heat kernels as the basis for our frame design. A heat kernel is defined on the graph spectrum as:

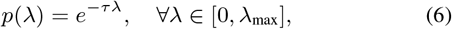

where *τ* is a free parameter that specifies the extent of lowpass filtering; the greater the *τ*, the narrower the lowpass band. Fig. 1 shows the heat SOSKS (HeatSOSKS) used in this study, consisting of *J* = 200 kernels. As *τ* → ∞, the kernel converges to a Kronecker delta in the spectral domain, which corresponds to a scaled version of the graph’s first Laplacian eigenvector in the vertex domain. On the other hand, as *τ* → 0, the kernel becomes a constant function in the spectral domain, i.e. an all-pass filter, corresponding to a Kronecker delta in the vertex domain. For each *τ*, we approximated *p*(*λ*) by a Chebyshev polynomial—thus approximating a minimax polynomial minimizing an upper bound on the approximation error [22]—and applied filtering as in (2); the polynomial order required to obtain a suitable approximation of the heat kernel varied based on the choice of *τ*, spanning the range 32 to 159 for *τ* = 5 and *τ* = 1000, respectively.

**Fig. 1.**
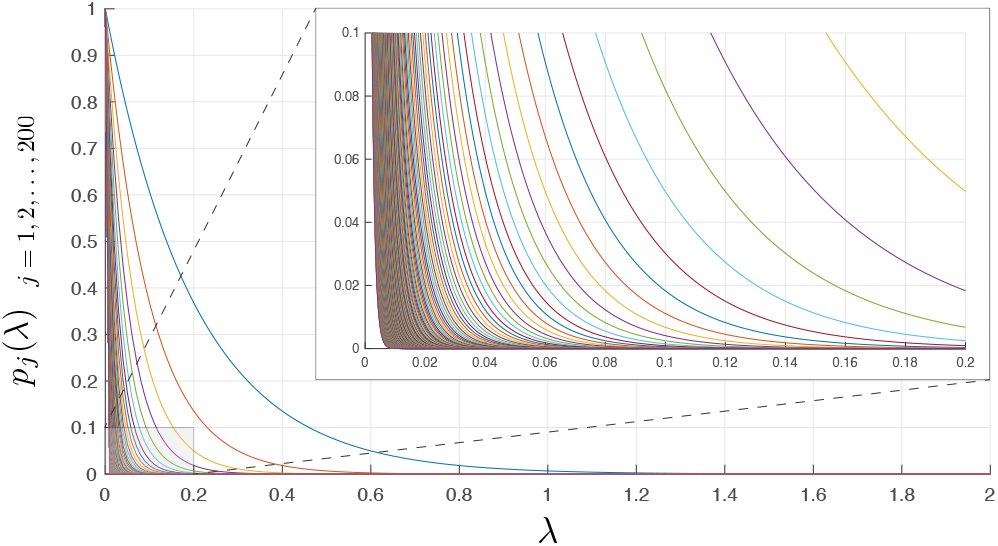
Heat system of spectral kernels (HeatSOSKS); *J* = 200 heat kernels for *τ* = 5 to 1000 with step of 5.

### 2.3. Dataset

We obtained structural MRI and resting-state fMRI data from the group of 100 unrelated subjects (54% female, mean age = 29.11 ± 3.67, age range = 22-36) of the Human Connectome Project (HCP) dataset [23]. We used the HCP minimally preprocessed data [24], that ensure the necessary co-registration accuracy to apply this method. Resting-state fMRI data for each subject consist of two sessions, each with 1190 time frames after excluding the first 10 to ensure fMRI signal stability. A thorough description of the image acquisition parameters and preprocessing steps can be found in [24].

### 2.4. Cerebral hemisphere cortex (CHC) graph

To characterize the convoluted structure of the cerebral cortex, we design subject-specific, voxel-resolution graphs that encode hemispheric morphology [14, 16]. At the core of the design is the use of a volumetric representation of a subject’s cerebral hemisphere cortex, i.e. the cortical ribbon, extracted from a T1-weighted MRI image with FreeSurfer [25]^1^. A graph vertex is then associated to each voxel that lies within the 3D ribbon. Graph edges are defined based on a two step procedure. First, an edge is associated to vertices for which the associated voxels are adjacents based on 26 neighbourhood connectivity in 3D [15]. Second, spurious edges derived from step one that are anatomically unjustifiable—e.g. edges that connect adjacent voxels that lie on the opposite sides of narrow sulci—are pruned out by using the pial and white surfaces. We thus obtain two graphs per subject, one per hemisphere, entailing a unique harmonic basis [14] using which fMRI data can be characterized [16, 17].

### 2.5. Structure-informed functional signatures from fMRI

For each subject, and each resting-state fMRI sessions, we extracted *P* static seed-based co-activation patterns (SSBCAP), each associated to one cortical region as defined by an atlas [18]; we studied parcellations with *P* = 200 and 400. Each SSBCAP was defined with the method introduced in [19]; by considering each region as a seed and averaging the top 15% of the frames (time-points) from the fMRI session that showed maximum amplitude in that region. Two graph signals were then extracted from each SSBCAP using the two CHC graphs, resulting in 2*P* graph signals per fMRI session. Each graph signal was *ℓ*_2_-normalized and decomposed using the Heat-SOSKS (*J* kernels), resulting in a total of *F* = 2*PJ* structure-informed functional signatures (SIFS) for each session.

### 2.6. Fingerprinting

We assessed the fingerprinting power of SIFS using a conventional procedure used for FC matrices [6, 26]. Let **y**_*s*,1_ ∈ ℝ^*F*^ and **y**_*s*,2_ ∈ ℝ^*F*^ denote the SIFS vector of subject *s* (*s* = 1, …, *S*) for sessions 1 and 2, respectively. An *S* × *S* identifiability matrix **M**^SIFS^ was defined, with elements 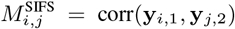, where corr denotes Pearson’s correlation coefficient. An identifiability measure *I*_diff_ ∈ [0, 100] was then derived as [6]: *I*_diff_ = (*I*_self_ −*I*_others_) × 100, where *I*_self_ and *I*_others_ denote the average of the main diagonal and off-diagonal elements of **M**, respectively. The higher the value of *I*_diff_, the higher the individual fingerprint overall across the subjects.

### 2.7. Feature ranking and selection

To be able to compare fingerprinting performances with the ones obtained from conventional FC, we reduced the feature space to retain at most an equal number of features as that provided by FC, i.e. the number of upper triangular elements in a FC matrix, (*P* ^2^ − *P*)*/*2. In order to rank the features, we used a measure inspired by the notion of differential power as given in [1]. Given that elements of **M**^SIFS^ are defined based on Pearson’s correlation between pairs of features from two sessions, the contribution of each feature to the resulting correlation can be treated as a measure of significance of that feature for fingerprinting, thus used to rank the features. Let 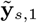 and 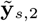 denote the z-scored versions of feature vectors **y**_*s*,1_ and **y**_*s*,2_, respectively. A measure of the significance of the *k*-th element in the feature vector can be defined as:

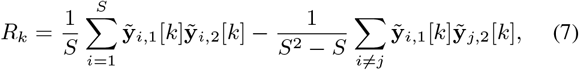

where *k* = 1, …, *F*. The larger the value of *R*_*k*_, the greater the impact of edge *k* overall along the population in fingerprinting. Having ranked the features, we selected different subsets of the top ranked features for assessing the fingerprinting power of SIFS; detailed selection choices given in Section 2.8.

### 2.8. Fingerprinting performance

To validate robustness of the fingerprinting performance as measured by *I*_diff_ relative to the choice of subjects, we performed a bootstrap analysis, consisting of four steps: (i) random selection of 80 subjects out of the 100, treated as the train set, whereas the remaining 20 subjects were treated as test set; (ii) feature selection as described in 2.7 using the train set; (iii) computing *I*_diff_ on the test set subjects using the feature categories selected in (ii); (iv) computing *I*_diff_ on the test set subjects using FC. Steps (i) to (iv) were then repeated 100 times, each time resulting in two *I*_diff_ values, one for SIFS and one for FC. Furthermore, to assess the effect of the number of features selected for SIFS in fingerprinting performance, we varied the number of selected features for SIFS relative to number of features in FC (number of upper triangular elements in FC graph); in particular, we validated the performance using 10% up to 100% features compared to that in FC, with a step of 10%.

## 3. RESULTS AND DISCUSSION

Fig. 2 shows the fingerprinting performances using SIFS vs FC. In particular, for SIFS, fingerprinting power was assessed using different sets of features selected based on the ranking procedure described in Section 2.7. Using both parcellation schemes, overall identifiability using SIFS outperforms that of using FC not only when using the same number of features as FC, but also less features down to 70%. Moreover, both FC and SIFS provide greater fingerprinting power when assessed on the parcellation with more brain regions. In the following, results on SIFS are presented for the setting where number of features is equal to that of FC.

**Fig. 2.**
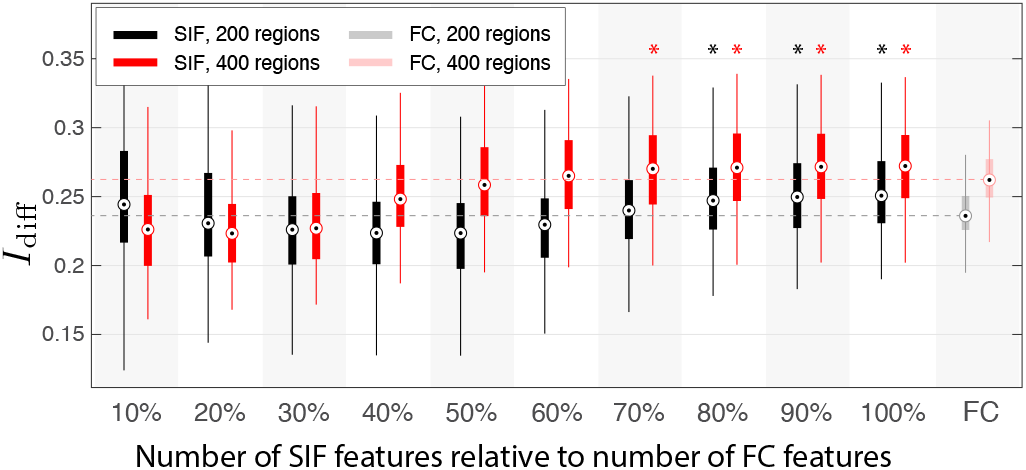
Fingerprinting performance using SIFS vs FC. Results are presented using two different cortical parcellations. An asterisk on top of a box-plot indicates SIFS performance being significantly greater than that of FC based on a paired t-test (p-value *<* 0.01).

Given that a subset of SIFS were selected based on the ranking procedure, we assessed how different kernels and different brain regions/networks were selected. Each brain region as provided by the atlas belong to one of the seven canonical functional brain networks [27]. Fig. 3 shows how the fraction of selected heat kernels varies between the different brain networks. In both parcellations, regions within the limbic network are more dominantly linked with heat kernels with low *τ* (less strict lowpass filters), whereas regions within the other networks are more dominantly linked to heat kernels with higher *τ* (more narrow-band lowpass filters). In particular, SSBCAPs associated to the visual and dorsal attention networks are more dominantly linked with high *τ* whereas SSBCAPs of regions within high-order cognitive networks such as the default mode and frontoparietal network provide features across a rather uniform range of *τ*.

**Fig. 3.**
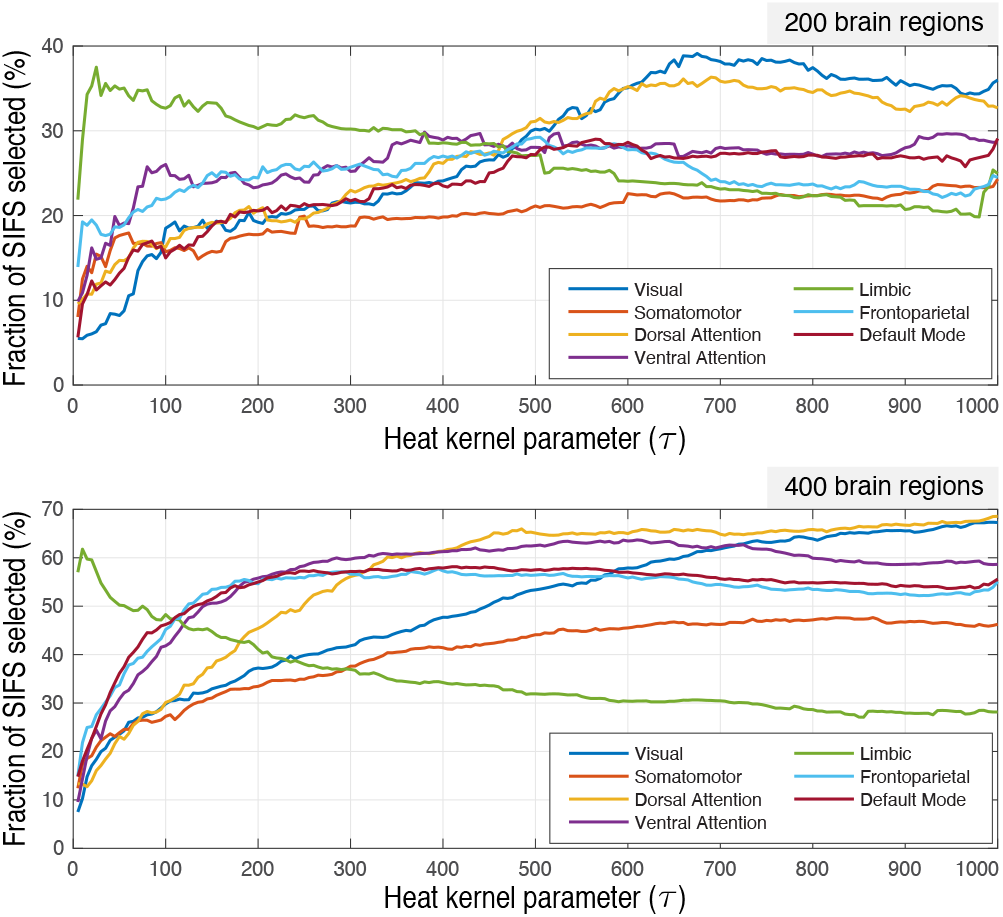
Significance of different heat kernels for fingerprinting in relation to different brain networks. The percentage values represent the fraction of times a kernel was selected for regions within a network out of the total possible number of possibilities; max possibility per network per kernel (i.e. a value of 100%) would correspond to: number of regions within network × number of bootstraps × two hemispheres.

Furthermore, it is insightful to observe that the number of SIFS selected per network notably differs depending on the parcellation resolutions; see Fig. 4. When using a finer cortical parcellation (400 regions), signatures are more dominantly selected from high-level cognitive networks—default mode, frontoparietal, dorsal and ventral attention networks; this observation corroborates findings presented in [28], which suggest that key information for differentiating subjects relies in high-level cognition regions, rather than in regions within the visual, somatomotor and the limbic networks. In contrast, when using a coarser cortical parcellation (200 regions), the distribution of selected features across the networks is more uniformly spread. This observation may explain the superior identifiability power of SIFS using 400 brain regions over 200, as shown in Fig. 2. To further scrutinize the distribution of selected features associated to each of the 400 regions, the latter were ranked based on their significance in fingerprinting, cf. Section 2.7. Resulting nodal significance measures were then ranked, such that a highly ranked region implies that SIFS associated to that region being more dominantly selected. The results are presented as a cortical projection in Fig. 5. Most notably, within the left hemisphere, high rank regions are observed in the lateral temporal lobe and medial frontal lobe, which encompass the default mode network, as well as several notable regions within the frontal lobe. Within the right hemisphere, sporadic highly ranked regions that fall within the ventral attention, default mode and frontoparietal networks are observed, as well as several regions within the dorsal attention network. Overall, these finding suggest the greater importance of signatures associated to high-level cognitive networks for fingerprinting.

**Fig. 4.**
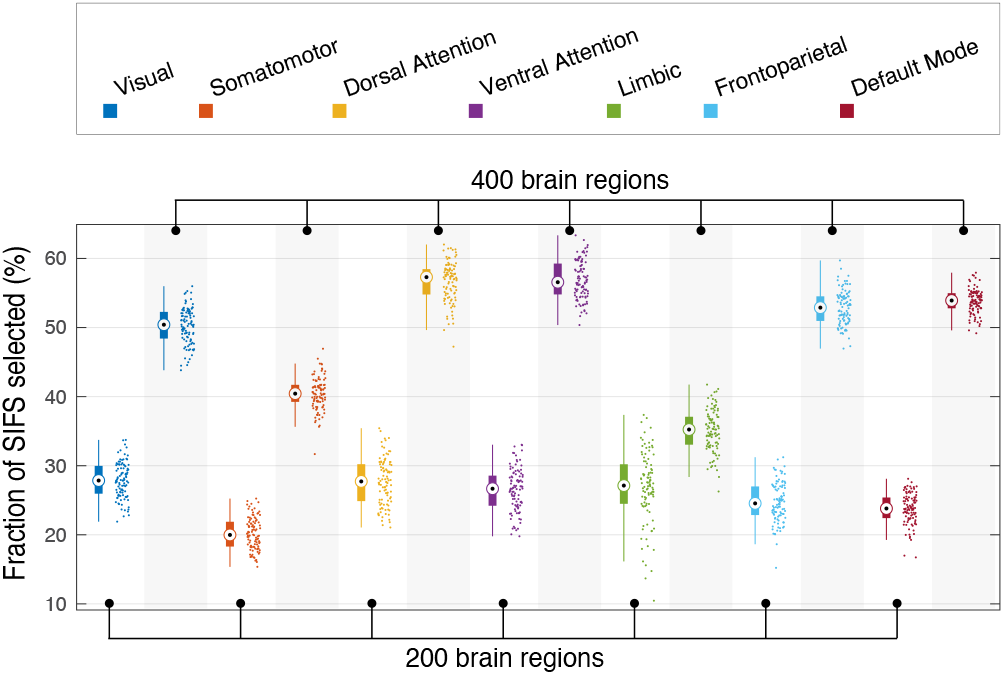
Percentage of selected features out of the total set of features belonging to specific networks; results over 100 bootstrap iterations, each dot showing result from a single bootstrap round.

**Fig. 5.**
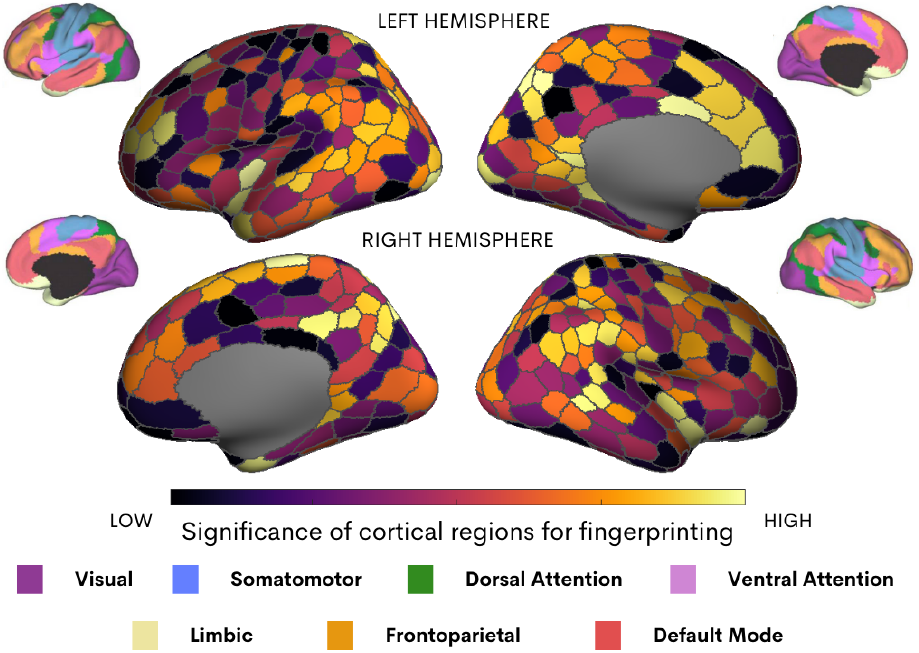
Significance of cortical regions in fingerprinting. Highest and lowest significance correspond to first and last ranking based on the procedure in Section 2.7. Each cortical projection is accompanied with the projection of the seven canonical functional networks [27].

## 4. CONCLUSIONS

We presented a method to extract structure-informed functional signatures from fMRI data using high resolution cortical graphs via graph signal processing principles. Results demonstrated that the proposed signatures provide greater overall subject identifiability than correlation-based functional connectivity. Moreover, the proposed method identified a specific cortical profile, showing high-level cognitive regions playing a greater role in subject identi-fication. Future work is needed to establish a more rigorous scheme for defining suitable graph frames [29], to explore the possibility of scaling up the feature space to derive voxel-specific signatures per graph kernel [30, 31], and to extend the methodology to incorporate white matter fiber architecture for more exquisite adaptation to underlying brain structure [13, 32].

## Footnotes

1 https://surfer.nmr.mgh.harvard.edu

